# Scale-sensitive Mouse Facial Expression Pipeline using a Surrogate Calibration Task

**DOI:** 10.1101/2024.05.15.594417

**Authors:** Andre Telfer, Oliver van Kaick, Alfonso Abizaid

## Abstract

Emotions are complex neuro-physiological states that influence behavior. While emotions have been instrumental to our survival, they are also closely associated with prevalent disorders such as depression and anxiety. The development of treatments for these disorders has relied on animal models, in particular, mice are often used in pre-clinical testing. To compare effects between treatment groups, researchers have increasingly used machine learning to help quantify behaviors associated with emotionality. Previous work has shown that computer vision can be used to detect facial expressions in mice. In this work, we create a novel dataset for depressive-like mouse facial expressions using varying LypoPolySaccharide (LPS) dosages and demonstrate that a machine learning model trained on this dataset was able to detect differences in magnitude via dosage amount.

## 1. Introduction

In preclinical research, animal models are often used in the study of emotions and to develop treatments with the goal of translating findings to humans. A popular animal for these studies is mice as they possess many physiological similarities with humans, are quick to produce, and have a well-mapped genome [3]. Being able to understand and accurately quantify emotions in mice, therefore, has significant implications for human mental health research and treatment. Unlike in human research, mice cannot self-report emotions; instead, emotionality is typically quantified through physiology or behavior. Physiological measures such as quantifying the increased presence of cortisol in stress studies are costly and invasive, which can impact later measurements. Behavioral data, on the other hand, can be captured passively and cheaply using video and used as a screen for drugs that stabilize affective states, like antidepressants.

The Open Field Test (OFT) is one popular behavioral screen to measure behaviors associated with anxiety and depressed motivation [2]. Another commonly used behavioral test is the Mouse Sickness Scale, where experimenters observe mice for features that indicate if the animal is unwell by observing such characteristics as ataxia (low locomotor activity), piloerection (hair sticking up), hunching, and ptosis (eyelids closing [6]). The Mouse Grimace Scale is designed to capture pain expressions and is in veterinary and preclinical settings [8]. One of the advantages to scales such as the Mouse Sickness Scale and Mouse Grimace Scale is that they can often be scored in the home cage, whereas behavioral paradigms require the mouse to be put in a separate testing environment, which can induce anxiety even if mice are acclimated repeatedly.

Behavior analysis has traditionally been a manual endeavor that is time-intensive and introduces noise and human-induced errors. Automated tools have become popular for quantifying emotionality. The rapid advancement of Deep Learning has led to open-source machine learning software that can outperform commercial tools on fundamental tasks [9, 11]. AI has previously been applied to facial expressions in restrained mice in order to distinguish between a wide range of facial expressions induced using different stimuli such as pleasure, fear, and disgust [4]. With freely moving mice, researchers have developed a deep learning pipeline that emulates human scoring of the Mouse Grimace Scale from video using a stochastic frame extraction technique [1]. Our study builds on the Mouse Grimace Scale pipeline [1] using a model that can score batches of sampled frames. In addition, we create a novel dataset with depressive-like symptoms using lipopolysaccharide (LPS) to induce a controlled sickness response known to promote anxiety-like and depressive-like behaviors such as appetite loss and social withdrawal in both humans and mouse models [7]. Our dataset is designed to test for scale sensitivity by including multiple LPS dosages measured at multiple time points. We demonstrate that our trained model produces scores that align with what would be expected from scale-sensitive scoring of facial expressions.

## 2. Methods

### 2.1. Setup

We used male and female adult C57BL/J6 mice (N=40 mice, 2-4 months old) obtained from Jackson Labs (Bar Harbor, Maine) and mouse colonies bred at Carleton University. Mice were housed in pairs before the experiment. They had free access to food and water throughout the experiment. All animals were kept under standard laboratory conditions with stable temperature (21 degrees C), humidity (15%-20%), and a 12-hour light/dark cycle with lights on at 07:00 AM. All procedures were approved by the Carleton University Animal Care Committee (AUP 111623) and adhered closely to the Canadian Council for Animal Care. The equipment and configuration were selected based on previous studies [1].

### 2.2. Procedure

Data were collected from 40 mice (male=20, female=20). Mice were assigned to one of 4 treatment groups: saline (male=5, female=5), 0.1mg/kg LPS (male=5, female=5), 0.5mg/kg LPS (male=5, female=5), and 1.0mg/kg LPS (male=5, female=5). LPS dosages were selected based on previous works where 1.0mg/kg would be considered high and 0.1mg/kg falls below the minimum dose typically used for depression-like models [13]. Mice were singly housed for at least 72 hours before filming. All injections consisted of a 0.2cc drug or vehicle. Recordings were taken at 5 time points: acclimation, baseline prior to injection, 1-hour post-injection, 2 hours post-injection, and 4 hours post-injection (Figure 3).

**Table 1.** Weak video labels for surrogate classification task.

|  | Pre | 1h Post | 2h Post | 4h Post |
| --- | --- | --- | --- | --- |
| Saline | 0 | . | . | 0 |
| LPS 0.1mg/kg | 0 | . | . | . |
| LPS 0.5mg/kg | 0 | . | . | . |
| LPS 1mg/kg | 0 | . | . | 1 |

The experiment consisted of two cohorts, one male and one female. For the injected groups (saline, 0.1mg/kg LPS, 0.5mg/kg LPS, and 1.0mg/kg LPS), four mice were filmed each day and assigned a different treatment group in random order. Filming each class followed best practice as it helped prevent models from using changing background information to make predictions.

On the day of the experiment, the mice were taken from the colony room and brought to a testing room in the morning and recorded as they acclimated in the transparent viewing box for 10 minutes. An hour later, this process was repeated and the mice were filmed to create the baseline videos. Following the baseline recordings, mice that were assigned saline or LPS treatments were taken out of the box and injected intraperitoneally with 0.2cc. An hour after the injection, the mice were placed back in the transparent viewing box and recorded again for 10 minutes. Recordings were taken again an hour later (2 hours post-injection), followed by two hours later (4 hours post-injection), for a total of 5 videos. The timings of these recordings were selected as depressive and anxiety-like behaviors in LPS mouse models, such as decreased social exploration and appetite, often peaking at 2-4 hours [7].

### 2.3. Pipeline

The collected were used to train a computer program that combined a deep learning approach based on ResNet [5] with a mean aggregation inspired by Deep Sets [14] (Figure 5). and an existing facial-feature extraction pipeline created in [1]. The pipeline periodically extracts frames from the video (we selected every 10th frame) and employs DeepLabCut[9] to first extract facial landmarks such as the eyes, ears, and nose of the mouse. Using these landmarks we discarded frames where neither side of the face is completely visible (i.e. for the left side: Nose, Left eye, Left Ear). To train the DeepLabCut model to identify the facial landmarks we took an iterative approach of manually labeling 100 frames, then training a model, using the model to automatically label several times more frames, and finally manually correcting frames and retraining the model. We repeated this until 5,000 frames had been labeled and the DeepLabCut validation error was reported as less than 3 pixels (e.g. on average DeepLabCut predictions were less than 3 pixels from the ground truth). We then extracted body part positions for all frames in every video using the trained model.

Further preprocessing was done by rotating frames such that the nose-eye axis was level, flipping frames where the mouse was facing right to match the left side, and translating them such that the eye was centered before finally cropping frames to 224x224 pixels (Figure 2). Finally, each training sample was made by randomly sampling several frames (n=5) from a video.

**Figure 1.**
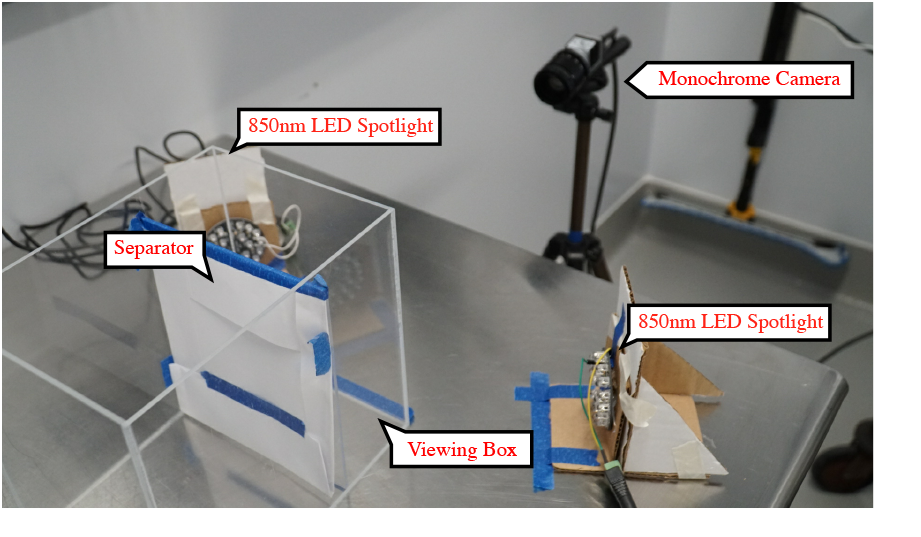
Example of filming layout.

**Figure 2.**
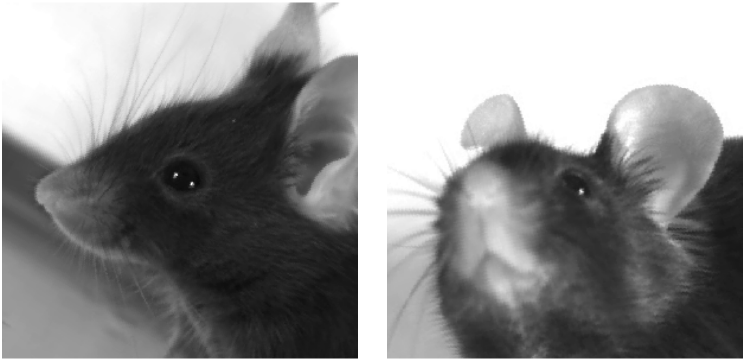
Example of a collection of preprocessed images from a single video, which would result in a single prediction from the model.

**Figure 3.**
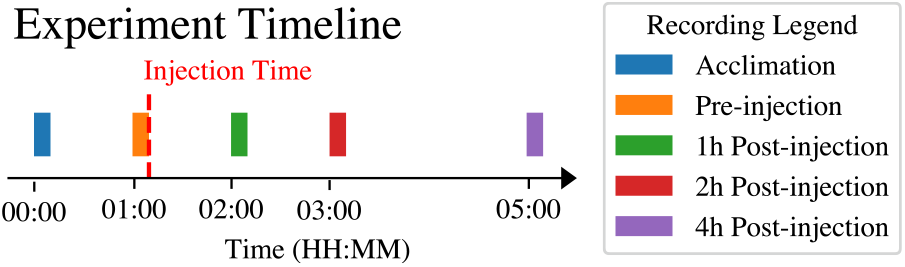
Timeline for recording videos (times are relative to experiment start time).

The collections of preprocessed frames were scored using a ResNet-based model that produced a single score from several frames using a mean aggregation inspired by Deep Sets [14] (Figure 5). The advantage of this approach is that: (1) the model can determine how much emphasis to put on each frame by assigning larger scores to frames that contain relevant information; (2) by using multiple frames the model has the advantage of having more information to make predictions; (3) loss is only calculated on the final prediction for the set of frames, rather than individual frames, greatly reducing the risk that the weak video label will not match any of the inputs and making training more consistent. Training was performed using kfolds cross-validation (k=10) with each validation split done according to day. In this way, mice were either used for training or evaluation (videos would never be split between both groups). The surrogate training task selected was for the model to classify high-dose LPS from the saline controls at Preinjection and 4 hours Post Injection 1. Final scores were produced by taking model predictions and subtracting the baseline pre-injection video for each mouse and normalizing all scores between [0, 1].

Following [1] we did not perform any image augmentation (e.g. additional rotation or masking) during training as empirical observations showed that did not improve performance. We used a standard cross entropy loss for our surrogate classification task and balanced the classes by generating more samples from the 1.0mg/kg LPS group. In total, 25000 training samples were generated for each class by sampling collections of frames from the same video. Details on learning rates, optimizer, and other hyper-parameters can be found on GitHub https://github.com/A-Telfer/mouse-facial-expressions-2023.

## 3. Results

Using posthoc Mann-Whitney U Tests to compare between groups at the same timepoint, the model indicated a strong ability to distinguish between saline/low-dose (0.1mg/kg) LPS groups and the high (1mg/kg) LPS group (p=0.0004 and 0.0003 respectively) at 4 hours post injections. The model could also clearly differentiate between Saline and High LPS groups much earlier at the 2-hour post-injection recording (p=0.0008). Differences between Low and Mid LPS doses were less pronounced and were not significant when using a Bonferroni correction for *α* = 0.05. The model was unable to differentiate between Low LPS and Saline groups. These results are consistent with previous literature examining anxiety and depressive-like behaviors associated with LPS which shows that these behaviors are most noticeable after 2 hours with dosages closer to our Mid/High groups [7].

### 3.1. 4 Hours Post Injection

Kruskal Wallis (p=0.0002)

| LPS | 0.1mg/kg | 0.5mg/kg | 1mg/kg |
| --- | --- | --- | --- |
| Saline | 0.6965 | 0.0080 | 0.0004 |
| LPS 0.1mg/kg |  | 0.0274 | 0.0003 |
| LPS 0.5mg/kg |  |  | 0.1779 |

### 3.2. 2 Hours Post Injection

Kruskal Wallis (p=0.0010)

3.2. 2 Hours Post Injection Kruskal Wallis ( $p=0.0010$ )
| LPS | 0.1mg/kg | 0.5mg/kg | 1mg/kg |
| --- | --- | --- | --- |
| Saline | 0.1011 | 0.0037 | 0.0008 |
| LPS 0.1mg/kg |  | 0.0592 | 0.0434 |
| LPS 0.5mg/kg |  |  | 0.4877 |

### 3.3. 1 Hour Post Injection

Kruskal Wallis (p=0.1973)

3.3. 1 Hour Post Injection Kruskal Wallis ( $p=0.1973$ )
| LPS | 0.1mg/kg | 0.5mg/kg | 1mg/kg |
| --- | --- | --- | --- |
| Saline | 0.6965 | 0.1530 | 0.4727 |
| LPS 0.1mg/kg |  | 0.0360 | 0.2743 |
| LPS 0.5mg/kg |  |  | 0.4379 |

### 3.4. Preinjection

Kruskal Wallis (p=0.2229)

3.4. Preinjection Kruskal Wallis ( $p=0.2229$ )
| LPS | 0.1mg/kg | 0.5mg/kg | 1mg/kg |
| --- | --- | --- | --- |
| Saline | 0.2031 | 0.7133 | 0.9097 |
| LPS 0.1mg/kg |  | 0.0483 | 0.0905 |
| LPS 0.5mg/kg |  |  | 1.0000 |

## 4. Discussion

In this paper, we developed a novel dataset for scoring depressive-like behaviors using LPS injections. We used a Deep Set [14] inspired approach with a surrogate classification task to allow the model to discard less valuable frames. We show that our final results are similar to those expected from a scale-sensitive model, with differentiation between dosage levels and over time (Figure 6). We validated the model by using occlusion-based explainability (Figure 7). For future directions we aim to test the model’s ability to translate to traditional paradigms for inducing stress-like responses such as Chronic Social Defeat.

**Figure 4.**
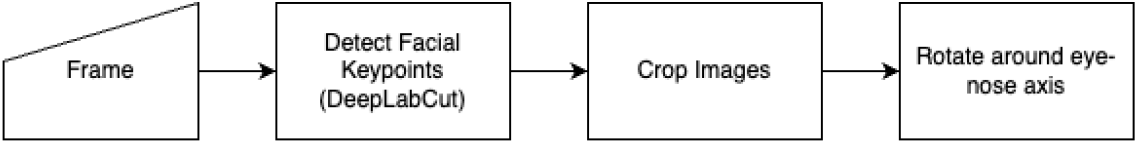
Preprocessing pipeline inspired by [1] with an additional step to rotate the frames such that the eye-nose axis aligns between frames, similar to traditional facial recognition preprocessing done for humans [12].

**Figure 5.**
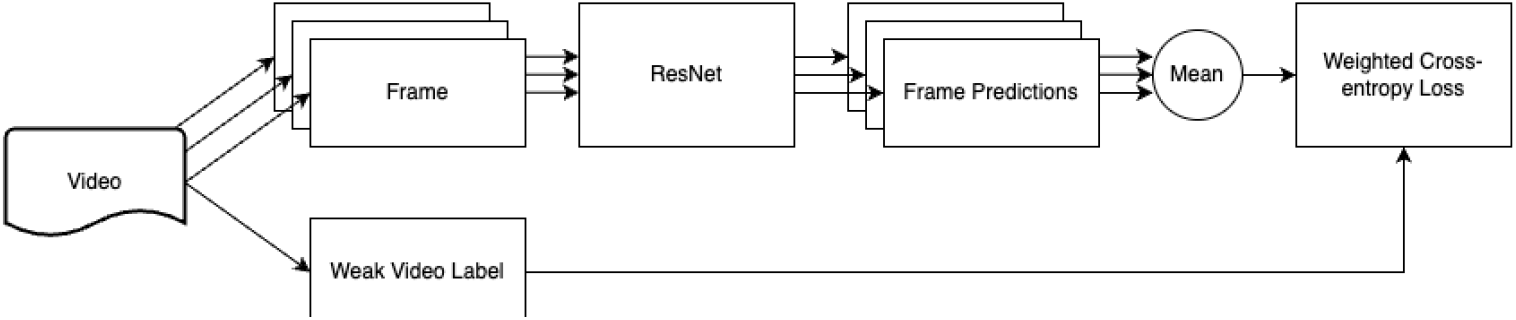
Model design for scores collections of frames from the same video. We used an aggregate method to combine scores from multiple frames inspired by Deep Sets [14] allowing the model to weigh how much each frame contributes to the final score (e.g. poor quality frames perhaps to activities such as grooming can be ignored without the need for further filtering steps).

**Figure 6.**
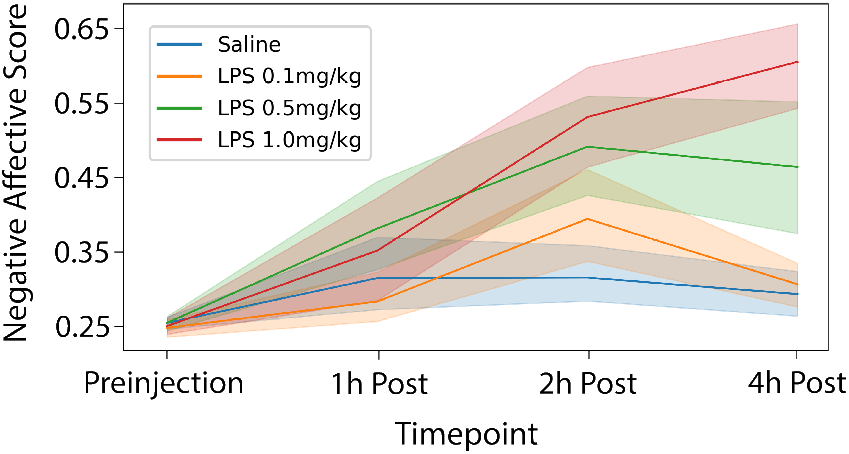
Plotting out the model scores reveals desirable characteristics of a scale-sensitive model, such as higher LPS doses having larger Negative Affect Scores.

**Figure 7.**
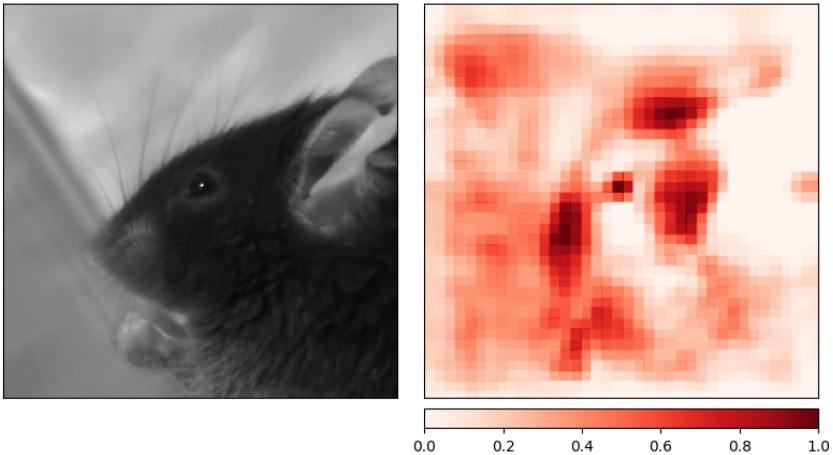
We used an Occlusion-based explainability technique that iteratively hides part of the image and observes its impact on the resulting class prediction [10]. This explainability method helps localize features important to the model, and we used it to verify that the model was using facial features in its predictions. Regions of interest, such as the cheeks and eyes, are also used in the Mouse Grimace Scale. We note that parts of the background also influenced the model score, suggesting that future models could benefit from a background-masking step.

## 5. Data Availability

The 200 captured videos and 5,000 annotated frames are available from the primary author.

